# Detecting Zero-Inflated Genes in Single-Cell Transcriptomics Data

**DOI:** 10.1101/794875

**Authors:** Oscar Clivio, Romain Lopez, Jeffrey Regier, Adam Gayoso, Michael I. Jordan, Nir Yosef

**Author notes:** Corresponding author. (N.Y.).

## Abstract

In single-cell RNA sequencing data, biological processes or technical factors may induce an overabundance of zero measurements. Existing probabilistic approaches to interpreting these data either model all genes as zero-inflated, or none. But the overabundance of zeros might be gene-specific. Hence, we propose the AutoZI model, which, for each gene, places a spike-and-slab prior on a mixture assignment between a negative binomial (NB) component and a zero-inflated negative binomial (ZINB) component. We approximate the posterior distribution under this model using variational inference, and employ Bayesian decision theory to decide whether each gene is zero-inflated. On simulated data, AutoZI outperforms the alternatives. On negative control data, AutoZI retrieves predictions consistent to a previous study on ERCC spike-ins and recovers similar results on control RNAs. Applied to several datasets and instances of the 10x Chromium protocol, AutoZI allows both biological and technical interpretations of zero-inflation. Finally, AutoZI’s decisions on mouse embyronic stem-cells suggest that zero-inflation might be due to transcriptional bursting.

## Introduction

In single-cell RNA sequencing (scRNA-seq) data, some genes have more zero measurements than can be modeled by a unimodal distribution centered at the mean expression level [1]. These zeros may be due to technical factors such as limited capture efficiency and sequencing depth, particularly for non-UMI protocols [2] and for infrequently expressed genes [1]. Zeros may also result from biological factors such as stochasticity inherent to the process of transcription (i.e, bursting) [3] or more stable differences between the milieus of genes that are present in different cell states.

Zero-inflated models are therefore commonly used for analyzing these data [1]. A first notable example is ZIFA [4], which uses a zero-inflated factor analysis to model the log-normalized data. ZINB-WaVE [5] and scVI [6] instead both rely on a zero-inflated negative binomial (ZINB) distribution to model the observed counts. Under the ZINB distribution, zeros can be attributed to either the limited sampling effects (NB) or to “surprising” zeros (ZI) which are not accounted for by NB. Indeed, it has previously demonstrated that under the scVI model, zeros attributable to the NB component better reflect the limitation in mRNA capture efficiency whereas the ZI component has a stronger association with the extent of read alignment errors [6].

A recent study [7] however questioned the universality of zero-inflation added to NB models for scRNA-seq data. This analysis was primarily based on datasets of “negative controls” (e.g., ERCC spike-ins) – namely exogenous transcripts that are constitutively expressed, thus reducing the contribution of biological factors as a cause for zero measurements. While the author found these negative controls to be adequately modeled by a NB with no zero-inflation, the question remains of whether this is also the case for cell-endogenous transcripts, and whether in such cases zero-inflation is a property specific only to a subset of the genes (e.g., due to their promoter kinetics [8]).

We address these questions by proposing a gene-specific treatment of zero-inflation. We present AutoZI, a novel generative model for scRNA-seq data which employs a spike-and-slab prior [9, 10] on a zero-inflation mixture assignment for every gene (Section 1). We propose a tractable inference procedure for AutoZI using variational methods (Section 2). A Bayesian decision rule based on AutoZI’s variational distribution gives a decision boundary between inferred ZINB and NB genes (Section 3). On simulated datasets, AutoZI outperforms other approaches in identifying zero-inflation (Section 4). On real datasets, AutoZI labels as zero-inflated only a small fraction of the negative controls (spike ins and control RNAs) while doing so for larger fractions of the endogenous genes, although this fraction tends to decrease with technical improvements in library preparation protocols (Section 5). In an application of AutoZI to mouse embryonic stem cells (mESC), we find that it is capable of distinguishing genes with a likely bursty promoter kinetics, labeling them as ZI. Such results suggest that : (i) negative binomial may not be globally appropriate for scRNA-seq data and that zero-inflation might be required to ensure a good fit for a significant fraction of the genes, and (ii) the patterns of genes detected as zero-inflated may be interpreted from both biological and technical perspectives.

## 1 The AutoZI probabilistic model

For each gene *g*, latent variable *δ*_*g*_ ~ Beta(*α, β*) indicates the probability of the absence of zero-inflation shared across all cells. Priors *α* and *β* are set to 0.5 to enforce sparsity while keeping symmetry. Latent variable *m*_*g*_ ~ Bernoulli(*δ*_*g*_) dictates whether gene *g* has its zero-inflation parameter sampled from the slab component (used for representing zero-inflation) or the spike component (otherwise) [9, 10]. For each cell *n*, let latent variable *z*_*n*_ ~ 𝒩 (0, *I*) be a low-dimensional random vector describing the cell’s biology, as in [6]. Let latent variable *l*_*n*_ ~ LogNormal 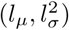) be a random scaling factor representing sequencing depth. For each gene *g* in each cell *n*, let latent variable 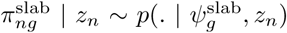 where 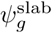 are parameters be a zero-inflation rate taking value in a set of non-negligible values (the “slab” component). Similarly, latent variable 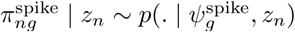 defines a zero-inflation rate taking value in a set of negligible values (the “spike”). Latent variable

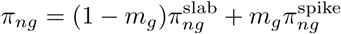

represents the effective zero-inflation rate. Finally, observed gene expression level *x*_*ng*_ is defined by

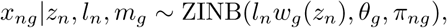

where *w* is a neural network taking value in the simplex (as in [6]) and *θ*_*g*_ are inverse-dispersion parameters learned via maximum likelihood.

In AutoZI, we define the spike and the slab by

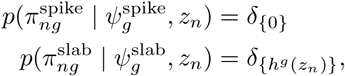

where *δ*_{*x*}_ denotes the Dirac distribution on *x* and *h* is a neural network taking values on the hypercube [τ_dropout_, 1]^*G*^. 0 < τ_dropout_ < 1 is used to lower bound the range of zero-inflation rates that can be accounted for by our model. Such a parameter avoids the possibility of having a nested mixture for the distribution *π*_*ng*_ (which happens for *h* = 0 and makes decision-making ill-defined). Without this constraint on the output of the neural network, we recover scVI if and only if the parameters *δ*_*g*_ are all equal to 0. AutoZI is yet another example of trade-off between interpretability and performance. Indeed, AutoZI is less flexible than scVI since it cannot attribute values to *π*_*ng*_ in the interval (0, τ_dropout_) but more interpretable since it provides a clear decision for gene-specific zero-inflation.

## 2 Variational inference

The marginal probability of the data *p*(*x*) is intractable. Therefore, we proceed to posterior approximation with variational inference in order to learn the model’s parameters. To approximate the posterior distribution, we first marginalize out the discrete random variables (*m*_*g*_)_*g∈G*_, as in collapsed variational inference. Each of the conditional distribution *p*(*x*_*ng*_ | *z*_*n*_, *l*_*n*_, *δ*) is a mixture of ZINB and NB distributions (with an identical NB component) with weight *δ*_*g*_. This makes the log-density *p*(*x*_*ng*_ | *z*_*n*_, *l*_*n*_, *δ*) tractable and differentiable with respect to the model’s parameters.

We approximate the posterior distribution of each {*δ*_*g*_, *z*_*n*_, *l*_*n*_}_*n∈N,g∈G*_ with a mean-field variational distribution:

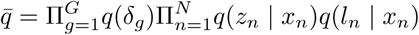

As in auto-encoding variational Bayes [11], each *q*(*z*_*n*_ | *x*_*n*_) follows a Gaussian distribution with a diagonal covariance matrix. Similarly, *q*(*l*_*n*_ | *x*_*n*_) follows a log-normal distribution. Parameters of these variational distribution are encoded via neural networks. For the global latent variables *δ*_*g*_, we use

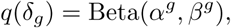

where each *α*^*g*^ and *β*^*g*^ are global parameters, numerically restricted to take values in (0, 1). We optimize the evidence lower bound (ELBO), derived as

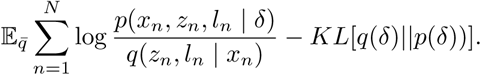

This objective function is amenable to stochastic optimization (as in [11]), which allows us to sample a fixed number of cells at each iteration [6] as well as from the variational distribution using the reparameterization trick [11] and its generalization to Beta distributions [12].

## 3 Detecting zero-inflation using Bayesian decision theory

Once the model fitted to data using the variational distribution we have at our disposal the variational posterior *q*(*δ*_*g*_) for each gene *g*. Our goal is to decide from this whether gene *g* is zero-inflated or not using discrete Bayesian decision theory [13]. Let 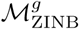 be the model for which *g* is zero-inflated. Similarly, let 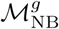 be the model for which *g* is not zero-inflated. Defining such hypotheses in scVI is not straightforward, especially at the gene-specific level. We can however rely on the latent variable *δ*_*g*_ of AutoZI to define formally zero-inflation by

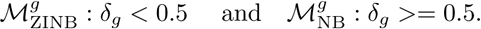

Let *K*_NB_ and *K*_ZINB_ be the costs of taking an inappropriate decision for an individual gene. We decide 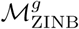 if and only if *q*(*δ*_*g*_ < 0.5) *> K*_ZINB_*/K*_NB_+ *K*_ZINB_. We note that this decision rule is approximate in the sense that we have only access to a variational approximation to the posterior. We focus in this paper on the case *K*_NB_ = *K*_ZINB_ = 1 for symmetry purposes. Under this particular setting and because *α* = *β*, our decision rule becomes equivalent to the Bayes factor of 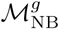 against 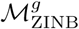 (a classical tool of Bayesian decision theory used in scVI [6] for differential expression).

## 4 Performance benchmarks on simulated datasets

We chose τ_dropout_ = 0.01 in all our experiments, as this value brought stable and balanced results on simulated datasets. As a consequence of AutoZI’s lower flexibility, we usually observed lower marginal log-likelihoods for AutoZI than for scVI. In this manuscript, we therefore focus on the performance of AutoZI at detecting zero-inflated genes. The ground truth for zero-inflation is unknown and not available for real datasets. Indeed, this is an open research topic. Consequently, we turned to simulations using Poisson log-normal distributions with added zero-inflation as well as Symsim [8], a realistic simulator for scRNA-seq data relying on Beta-Poisson distributions.

For both simulation frameworks, we benchmarked AutoZI against two decision rules based on maximum likelihood estimations: `Uni-MLE-Pi` and `Uni-MLE-LRT`. The former fits a univariate ZINB distribution to all genes and uses the fitted zero-inflation weight based on an arbitrary threshold to make a decision. The latter fits both a univariate ZINB and NB distribution and uses the differences in marginal likelihood Δ = LL^ZINB^ − LL^NB^ to perform decision. Examples of thresholds include AIC (resp. BIC) for selecting ZINB if and only if Δ ≥ 1 (resp. Δ ≥ ^log(*N*)/2^, where *N* is the size of the dataset). For AutoZI, we used the default hyperparameters from the scVI model and used the natural decision threshold at 0.5, denoted as “AutoZI default”.

### Poisson log-normal datasets

Each synthetic dataset contains 12,000 cells, 50 genes, and two cell types. Each cell has its gene expression drawn from a Poisson log-normal distribution whose mean and covariance matrix depend only on the cell type. A fraction *λ* of the genes are selected uniformly to be applied a Bernoulli mask with probability 0.1. We aggregate the results for *λ* ∈ {0, 0.25, 0.5, 0.75, 1} and report the ROC curves in Figure 1.

**Figure 1:**
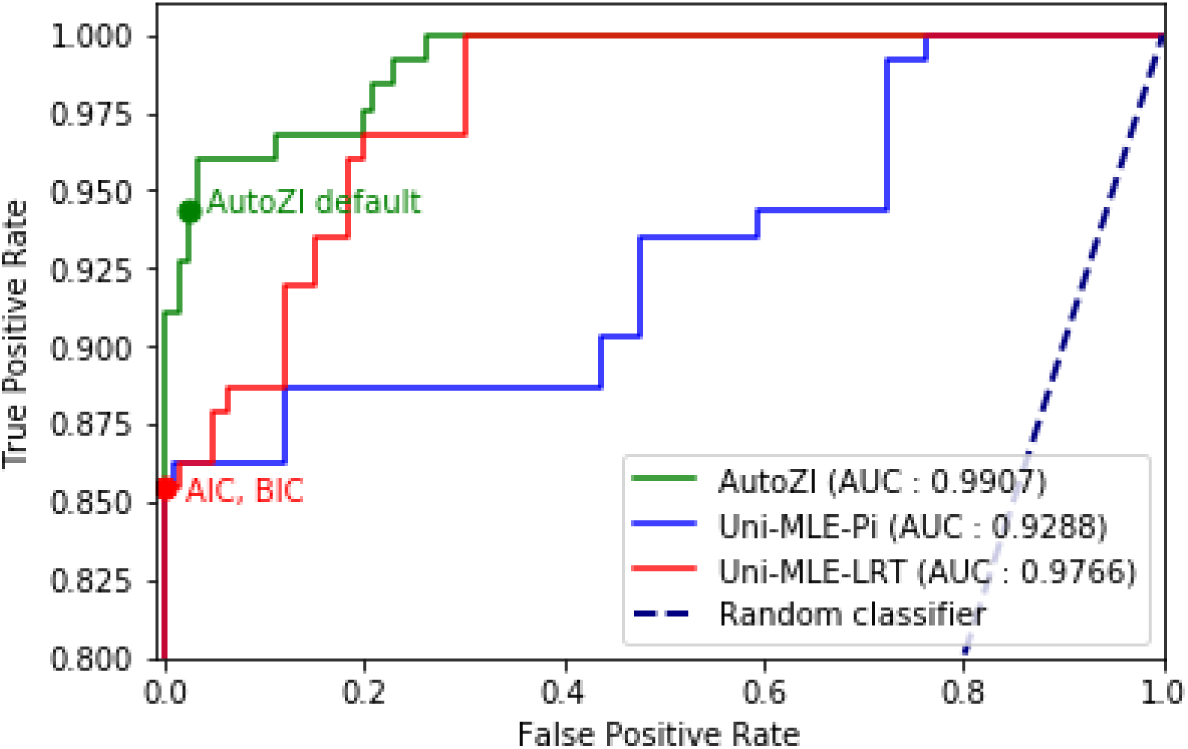
ROC curves on the aggregated Poisson log-normal dataset, with points corresponding to the default AutoZI decision rule, AIC and BIC.

The AutoZI model has a near-one AUC and `Uni-MLE-LRT` has a similar albeit slightly lower performance. In particular, AutoZI’s default decision rule leads to true positive (ZINB) and negative (NB) rates both superior to 94% whereas the AIC and the BIC do not yield false positives but their true positive rate is lower (85.5%), showing a bias in favor of the NB distribution. The `Uni-MLE-Pi` baseline procedure performs worse than these two models.

### Beta-Poisson datasets

For a more biologically relevant simulation framework, we used known kinetic models of stochastic gene expression such as the Beta-Poisson model. SymSim [8] provides a natural way of sampling data from such models and adding technical noise. SymSim first randomly samples the promoter on rate (*k*_on_), off rate (*k*_off_) and synthesis rate (*s*) for each gene, and then generates simulated “true” counts using a Beta-Poisson distribution. As a simplification, we label the genes with kinetic parameters *k*_on_, *k*_off_ < 1 as bimodal, *k*_on_ *>* 1 as unimodal with a non-zero mode (UNZ) and the rest as unimodal zero (UZ). The real regimes might depend additionally on *s* and *k*_on_*/k*_off_ but are negligible as a first analysis. True counts are then converted to observed counts by simulating processes such as capture, amplification and fragmentation (with UMIs in these datasets). We create a dataset of 100 genes with a bimodal distribution and another dataset of 100 genes with a UNZ distribution. Both datasets are subsampled multiple times (*n* bimodal genes and 100 − *n* UNZ genes, for *n* ∈ {0, 25, 50, 75, 100}) to create a sequence of datasets of 100 genes and 3,000 cells from a single cell type. We expect UNZ genes to be non zero-inflated, as these can be easily modeled by a NB distribution. Conversely, we expect genes that are bimodal in their true counts to be zero-inflated in their observed counts, due to limited sensitivity. For simplicity, we did not focus on UZ genes since their ground-truth category is not clear. We aggregate the results and report the ROC curves in Figure 2.

**Figure 2:**
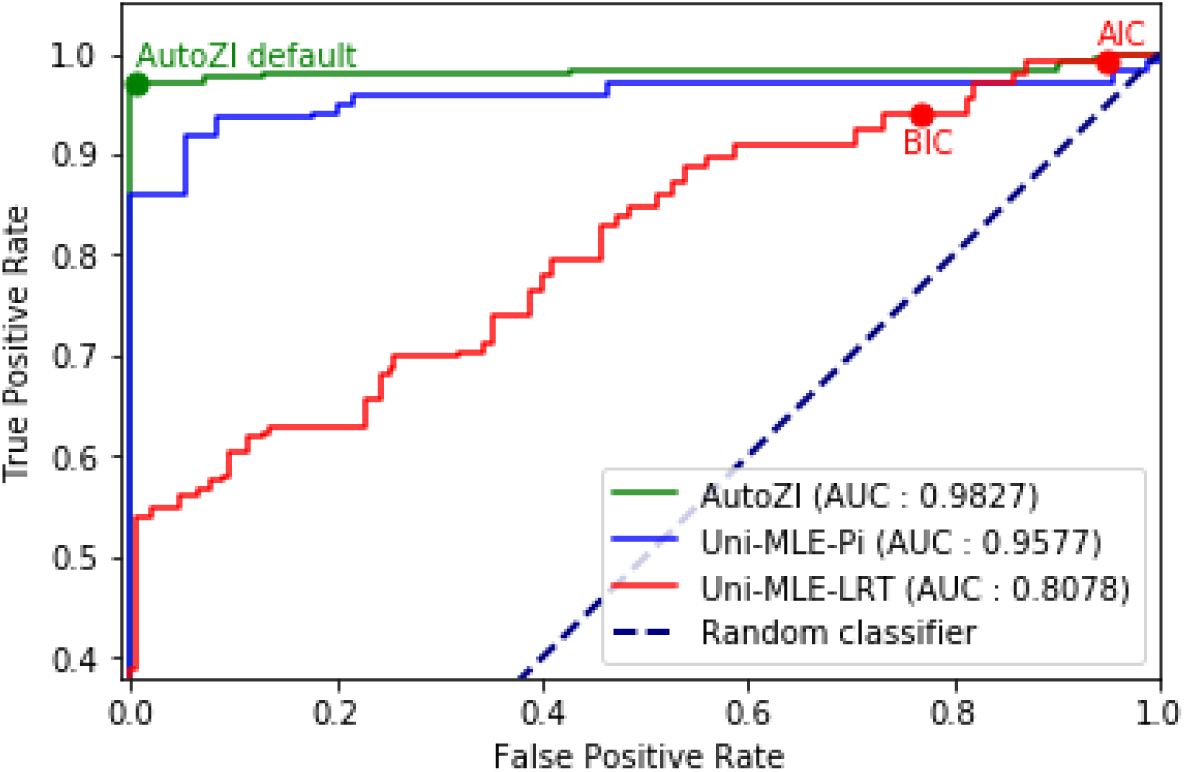
ROC curves on the aggregated SymSim dataset, with points corresponding to the default AutoZI decision rule, AIC and BIC.

AutoZI has the best area under the curve and its default decision rule classifies more than 96% of bimodal genes as ZINB and more than 99% of the UNZ genes as NB. Such results show that zero-inflation in our model might be suited to distinguish regimes of bimodality for kinetics of gene expression. Conversely, the `Uni-MLE-LRT` baseline hardly distinguishes the bimodal and UNZ regimes. In particular, both AIC and BIC are biased towards ZINB predictions, and thus poorly suited for zero-inflation analysis of real data.

### Robustness for lowly-expressed genes

An important consideration is the robustness of AutoZI’s performance for lowly expressed genes. Using Poisson-log-normal datasets with average expressions spanning from 10^*−*4^ to 10^2^, we found that AutoZI’s decisions became biased towards ZINB for genes with low average expression or, equivalently, average predicted negative binomial mean: notably, 92.7 (resp. 2.4%) of NB genes with average expressions above (resp. below) 1 were correctly retrieved. Such results suggest that the statistical problem of detecting zero-inflation becomes harder for low values of the negative binomial mean. Hence, for the following biological datasets, we train AutoZI on all genes but study its predictions only for those with average expression above 1.

## 5 Application to detecting zero-inflation in real datasets

A significant application is to investigate how zero-inflation affects real datasets and whether the decisions taken by AutoZI have a biological meaning.

### Negative control (ERCC spike-in and RNA)

We apply AutoZI to four droplet-based negative control datasets, spanning a wide range of experimental protocols (10x Chromium v1 [14], inDrops [15] and GemCode [16]) and based on ERCC spike-ins and control RNAs. Such datasets do not capture any biological process and the ERCCs were shown not to be zero inflated in [7]. To investigate whether AutoZI decision-making mechanism can reproduce similar results, we selected ERCC spike-ins and added the 100 most expressed control RNAs in each dataset for joint analysis. 32.4% of ERCC spike-ins and 100% of control RNAs had sufficient average expression. Hence we analyze between 24 and 44 spike-ins per dataset. We report decisions from AutoZI in Table 1. AutoZI has a perfectly symmetric prior and is therefore not biased towards any specific decision. However, 98.3% of ERCC spike-ins and 94% of the control RNA under study are retrieved as NB. These results corroborates the hypothesis from [7] that droplet-based ERCC spike-ins measurements are not zero-inflated and even may extend it to the control RNAs part of these datasets.

**Table 1:** Percentages of ERCC spike-ins and control RNAs predicted as ZINB by AutoZI on negative control datasets.

|  | Svensson et al. (1) [14] | Svensson et al. (2) [14] | Klein et al. [15] | Zheng et al. [16] |
| --- | --- | --- | --- | --- |
| ERCC | 0.0 | 8.3 | 0.0 | 0.0 |
| RNA | 4.0 | 14.0 | 0.0 | / |

### A collection of 10X biological datasets

We now check whether the fraction of ZINB genes in PBMC datasets sequenced using 10X Chromium is higher than in previous control RNA data and whether it decreases with the version of the protocol, as technical improvements may suggest. We focus on pbmc3k (10X v1.1.0), pbmc8k (10X v2.1.0) and pbmc10k (10X v3.0.0, with protein expressions). Their cell types were estimated using clustering techniques and marker genes or proteins. We focus on B cells, CD14+ monocytes, CD4 T cells and CD8 T cells. For training, we select genes both among the 1,000 most variable and expressed across all cell types in all datasets. For analysis, in each cell type, we select genes with sufficient average expression in all datasets, yielding between 203 and 228 genes per cell type. We report the percentages of ZINB genes predicted for each cell type by a gene-cell-type extension of AutoZI in Table 2. We note a higher general fraction of zero-inflated biological gene-cell-types (29.3%) than control RNAs (6%), indicating potential biological phenomena. However, for all cell types, we find that the fraction of predicted ZINB genes decreases with the version of 10X Chromium, potentially showing additional technical factors in zero-inflation. This may lead to a less straightforward conclusion than in [7] where droplet-based zero-inflation is stated as only biological.

**Table 2:**
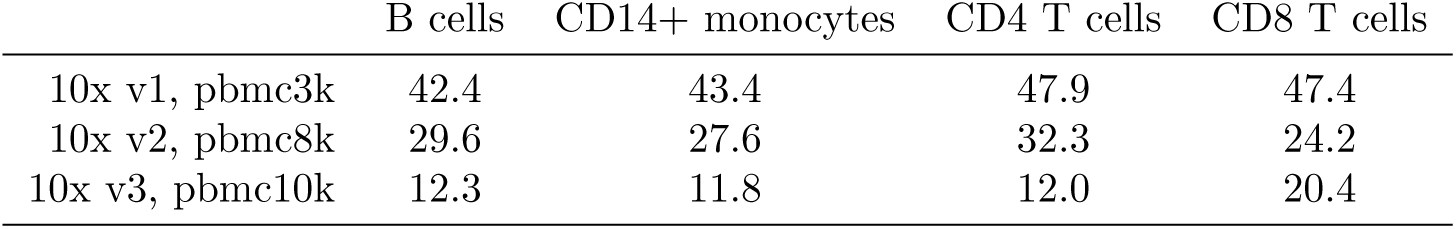
Percentages of genes predicted as ZINB by AutoZI on 10x PBMC datasets.

|  | B cells | CD14+ monocytes | CD4 T cells | CD8 T cells |
| --- | --- | --- | --- | --- |
| 10x v1, pbmc3k | 42.4 | 43.4 | 47.9 | 47.4 |
| 10x v2, pbmc8k | 29.6 | 27.6 | 32.3 | 24.2 |
| 10x v3, pbmc10k | 12.3 | 11.8 | 12.0 | 20.4 |

### Transcriptional burst kinetics of embryonic stem-cells

We previously explored the ability of AutoZI to capture bimodal genes from simulated Beta-Poisson models. Such model was fitted to an allele-specific scRNA-seq dataset of mESCs for characterizing transcriptional bursting [3] through estimates of the kinetic parameters *k*_on_, *k*_off_ and *s* for each gene. We investigated whether the decisions made by AutoZI could recapitulate the different regimes for the Beta-Poisson distribution (Bimodal, UZ, UNZ) in this dataset. Given the limited number of cells (188), we applied AutoZI to a larger mouse hybrid ESC dataset [17] (704 cells, one common allele), which was not used for the inference of the kinetic parameters, but should be biologically similar. After intersecting the gene lists in the two datasets, we further randomly filtered the genes and kept 52 bimodal, 52 UNZ and 29 UZ genes. Only one of the UNZ genes did not have sufficient expression for analysis. We report the decisions from AutoZI with respect to the kinetic parameters *k*_on_ and *k*_off_ in Figure 3. AutoZI predicts that 38 out of the 52 bimodal genes are zero-inflated and that 37 out of 51 UNZ genes are not zero-inflated. Taken together, our results suggest that AutoZI’s predictions of zero-inflation can be used not only to account for technical factors (which are evident by the decrease in ZI with progress in technology in Table 2) but may also reflect biological factors such as transcriptional bursting.

**Figure 3:**
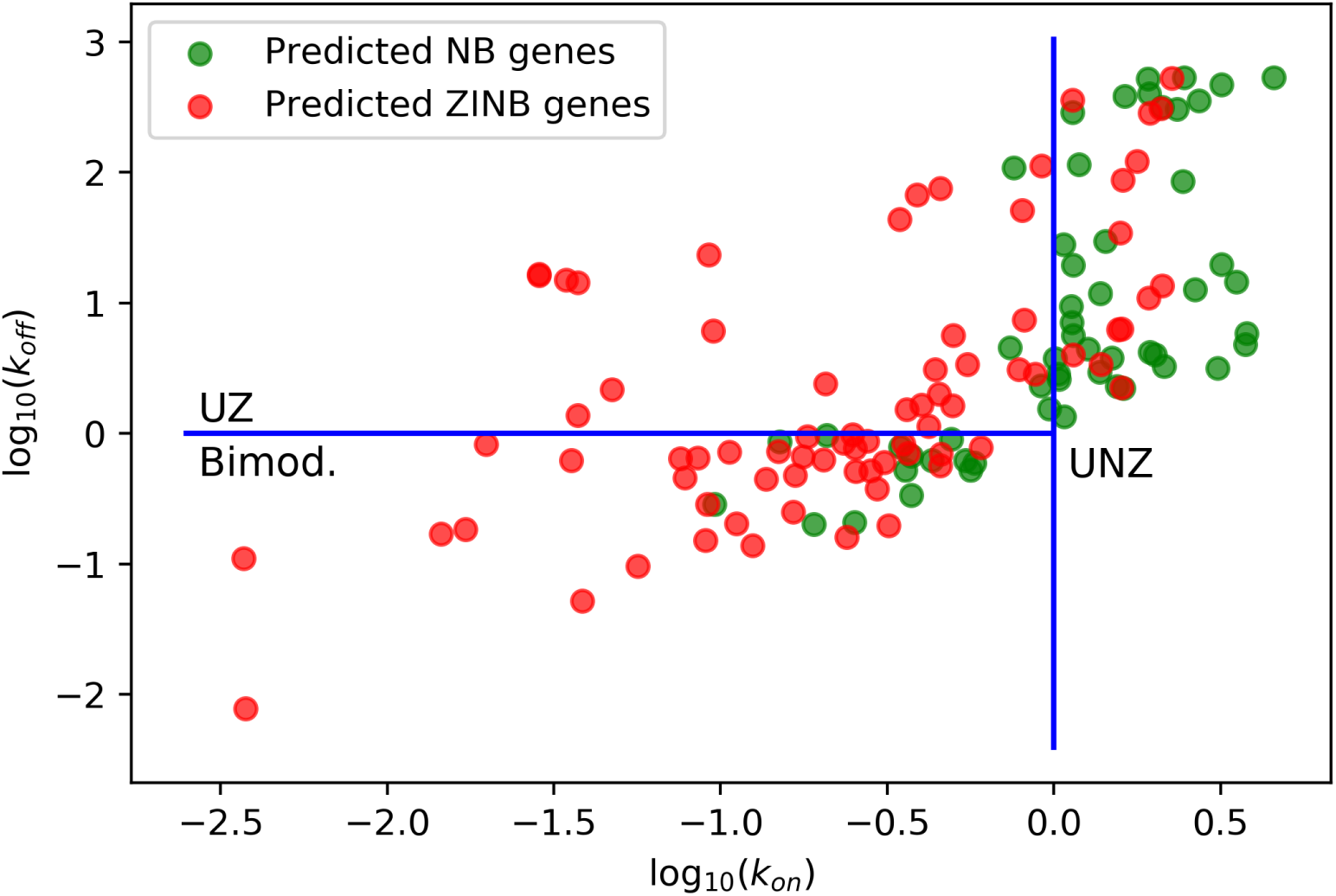
Genes from the ESC dataset from [17] plotted in the space of (*k*_on_, *k*_off_) kinetic parameters, with their NB/ZINB labels from AutoZI.

## Code availability

The implementation to reproduce the experiments of this paper is available at https://github.com/oscarclivio/AutoZI_reproducibility. The reference implementation of AutoZI is available at https://github.com/YosefLab/scVI.

## Acknowledgements

We thank Valentine Svensson whose work on zero-inflation in droplet scRNA-seq inspired the project, Chenling Xu for her help on ESC and 10X Genomics data and Pierre Boyeau for the cooperation on Poisson log-normal datasets. We also thank Matthew Jones and Zoë Steier for insightful biological discussions. Nir Yosef is a Chan Zuckerberg Biohub investigator.

